# Too hot for the weeds? Exploring the impact of climate change in herbaceous Convolvulaceae in Cerrado and Mata Atlântica biomes (SE Brazil)

**DOI:** 10.1101/2024.11.06.621913

**Authors:** Juliana Cruz Jardim Barbosa, Fábio Vitalino Santos Alves, André Luiz Costa Moreira, Benoît Loeuille, Lars W. Chatrou, Rosângela Simão-Bianchini, Ana Rita Giraldes Simões

## Abstract

**Aim:** We investigated the potential resilience to climatic change of nine species of weeds in Convolvulaceae, using predictive spatial modeling across two contrasting biomes in Southeastern Brazil: Cerrado (=Savanna) and Mata Atlântica (=Atlantic Forest). We inferred future changes in distribution area, the climatic variables that will be most impactful, and the potential occurrence of future climatic refuges.

**Location:** Southeastern Brazil (São Paulo, Brazil).

**Methods:** A total of 195, taxonomically vetted, distribution records were compiled for nine species of Convolvulaceae. Potential distribution areas were modelled in RStudio 1.3.1056 with R 3.6.3 using modleR; environmental layers used were the 19 bioclimatic variables with 30 seconds resolution. After a correlation analysis, four bioclimatic variables were selected for distribution modelling: Temperature Seasonality (BIO4), Mean Temperature of Wettest Quarter (BIO8), Precipitation of Wettest Month (BIO13) and Precipitation of Driest Quarter (BIO17). The species distribution modelling was performed using the Maxent algorithm for the present time (1960-1990) and for future projections (2050 and 2070), under two different scenarios of projections, moderate and pessimistic.. The final distribution models were generated through the selection of primary models with a minimum level of TSS (True Skills Statistics) equal to 0.7.

**Results:** The analysed species demonstrated different levels of response to climatic change: *Distimake aegyptius, D. dissectus*, and *Evolvulus pteurocaulon* exhibited a gain in the climatic suitability range, regardless of the future scenarios. Other species, such as *E. glomeratus, Ipomoea bonariensis*, and *I. alba* showed a decline in climatic suitability range, more accentuated in the pessimistic scenario. Three species showed a positive response in a moderate future scenario, but a decline in the most extreme projection. Areas that may act, in the future, as climatic refuges for the displaced species are also highlighted, in view of being prioritised for conservation.

**Main Conclusion:** This study reveals that climate change will have varied impacts on herbaceous species of Convolvulaceae occurring in the Brazilian Cerrado and Mata Atlântica biomes. Some species are likely to benefit from climate change, showing an increase in climatic suitability in both moderate and pessimistic future scenarios. Other species, however, are expected to experience a decline in suitable climatic conditions, in particular under the most pessimistic climatic scenario, and are more likely to be threatened in the future, due to the constriction of suitable habitats. The analysis also highlights that certain regions may serve as important climatic refuges for these species in the future. These findings emphasize the necessity for targeted conservation plans to protect the biodiversity of the Cerrado and Mata Atlântica phytogeographic domains and weed management strategies - not only in the present but also in future climatic conditions.

## 1. INTRODUCTION

According to the Brazilian deforestation monitoring, Brazil experienced an 11.6% reduction in deforestation rates in 2023 compared to 2022, with an estimated 1.83 million hectares of native vegetation lost. This marks the first annual decline since 2019, with the Cerrado biome accounting for 61% of the total deforested area, surpassing the Amazon’s share of 25% (MapBiomas, 2024). Brazil has six phytogeographic domains, characterised by Amazônia, Mata Atlântica, Cerrado, Caatinga, Pantanal and Pampa (Ab’Sáber, 2003), all suffering the impact of human occupation and climatic change in their levels of biodiversity.

Cerrado is the second largest phytogeographic domain in Brazil, surpassed only by the Amazon Forest. It accounts for 30% of Brazilian biodiversity, has also been fast declining. In 2009, its deforestation rate (0.32%) was more than double that observed in the Amazon (0.14%) in the same year. Yet, only a very small portion of its area is protected (Françoso, 2015): public protected areas cover only 7.5% of Cerrado and under Brazil’s Forest Code, only 20% of private lands are required to be set aside for conservation. (Strassburg, 2017).

The Atlantic Forest has suffered severe human impacts, resulting in the destruction of most of the primary coastal vegetation and transforming the landscape into areas of secondary vegetation (Püttker, 2008). Studies indicate that only 12.4% of the forest remains preserved. The reduction in the areas of native vegetation is mostly related to human actions. (Martins et, al. 2021). Recognized as a biodiversity hotspot with >18,000 species of plants (Flora e Funga do Brasil, 2023) and 3500 species of vertebrates (Figueiredo et al., 2021).’

Over the past 30 years, the Atlantic Forest has been a global priority for biodiversity conservation, being one of the top five Biodiversity Hotspots on the planet (Marques, 2020). Conservation units have been created to protect these areas, but studies indicate that even if all current Atlantic Forest remnants are protected, species could still be lost without appropriate ecosystem management and restoration efforts (Montovani, 1988; Zwiener et al., 2017).

In the state of São Paulo (SE Brazil), Cerrado and Atlantic Forest are the dominating phytogeographic domains. However, only 13.4% of the total surface is still covered by natural vegetation, corresponding to 3,340.774 hectares (Kronka, 2005). In São Paulo, Cerrado vegetation has been drastically reduced to only 0.81% of the state’s vegetation cover (Kronka et al., 2005), from its original 14% cover (São Paulo, 1997). Thus, the current vegetation of Cerrado in the state of São Paulo, corresponding to less than 7% of its original cover, has been further divided into thousands of small areas, surrounded and distributed in agricultural, reforestation and urban areas (Durigan et al., 2007).

In addition to drastic reduction in vegetation cover, growing urbanisation also induces localised changes in climatic conditions. The capital of the state, São Paulo, is the largest city in South America, with over 11 million residents. Since the colonization by the Portuguese, the state of São Paulo has been seriously affected by vegetation exploitation. In the 19th century, with the expansion of coffee cultivation, most of the original vegetation cover was devastated, from Serra da Cantareira, to the end of Serra do Mar (Secretaria do Meio Ambiente - SMA, 2008). In addition to the history of intense agricultural, hydroelectric reservoirs and industrial activity in the state of São Paulo, extreme urbanization has led to expanding residential construction, deforestation of large areas and the colossal increase of automobiles. Predictions show a very grim scenario for the native vegetation in a near future, and the potential of climate change and habitat reduction to drive extinctions of tropical plant species (Faleiro et al., 2013; Zwiener et al., 2017). In São Paulo, the temperature rise is already noticeable, with abnormally hot days in the winter and excessive rainfall in the summer, starting to interfere with ecological processes, the life cycle of species, and eventually leading to the extinction of several representatives of local flora and fauna (Marengo 2006).

Worldwide, climate change intrinsically affects species distributions, and in the future, many plant species will have to respond to rapidly changing abiotic conditions to be able to thrive (Thomas et al., 2004; Thuiller et al., 2005; McGaughran et al., 2021). In the 20th century, an increase of 0.65 °C in the average global temperature was recorded, more accentuated during the 1990s, with an increase of 0.2% to 0.3% in precipitation in tropical regions (Field et al., 2014). The resulting changes in temperature and precipitation regimes have already led species to alter their physiology, as a strategy to tolerate the warmer or drier conditions, shifting their distribution ranges to follow appropriate climatic conditions in space, as well as adapting some of their crucial life cycle events to favourable weather periods (Pecl et al., 2017). If species fail to adopt any of these strategies, species may face extinction (Román-Palacios & Wiens, 2020). A predicted increase in global temperature from 2ºC to 3 ºC may be devastating and can cause the transformation of forests to non-forest systems (Silvério et al., 2013; Seidl et al., 2016).

The responses of vegetation to climate vary considerably with the different spatial patterns and the effects of “time-lag”, important mechanisms dealing with the interactive effects of climate-vegetation (Wu et al., 2015). Extensive studies, focused on large-scale interactions between vegetation and climate, use simultaneous meteorological and vegetation indicators to develop models; however, the effects of elapsed time are less considered, increasing the uncertainties about these effects (Wu et al., 2015).

Predictive distribution modelling quantifies the ecological niche of species using known occurrence points and environmental data (Guisan & Zimmermann, 2000). This tool is crucial in ecology and conservation studies, including the expansion of invasive species (Peterson et al., 2007). The use of modelling to study the effects of climate change in the distribution of species is of great relevance because, in addition to helping to understand potential species distribution, it is a resource currently used to contribute to conservation work, showing the possible consequences of long-term climate changes and enabling the analysis of their effects on the distribution of organisms, measuring these changes in biodiversity and phytogeographic domains. Thus, the sustainable management of weeds is increasingly challenging due to the significant impact of climate change on herbaceous communities and, for example, their interactions with crops, which has a direct impact in agriculture. This emphasizes the need to develop effective and sustainable weed management practices that consider the simultaneous and complex effects of multiple climatic factors (Anwar et al., 2021).

For this study, we have selected a model group of nine herbaceous species in Convolvulaceae, a plant family which includes morning glories, bindweeds, dodders and the crop sweet potato. “Bindweeds” (*Convolvulus*, 203 spp) and “morning glories” (*Ipomoea*, 635 spp) together constitute nearly half of the species diversity of the family. Convolvulaceae, in general, are mostly herbaceous in habit, either erect, prostrate or climbing, and more rarely shrubs, sub-shrubs or trees (Staples, 2012; Simões et al. in prep.). Many are very successful in occupying disturbed or heavily urbanised environments, covering native vegetation or cultivated fields, for which they are often flagged as “weeds”, with a negative perception of invasive behaviour, which is, nonetheless, not always the case.

Brazil is an important centre of species diversity of Convolvulaceae, housing 426 of the 1,955 species distributed worldwide, of which 196 are endemic to Brazil. The most represented genera are *Ipomoea, Evolvulus, Jacquemontia* and *Distimake*, with the Cerrado domain concentrating the largest species diversity of these genera (Simão-Bianchini et al. 2020; Silva et al. 2018). In the state of São Paulo (SE Brazil), 116 species of Convolvulaceae are recorded, of which one, *Distimake hoehnei* (Petrongari & Sim.-Bianch.) Petrongari & Sim.-Bianch. is endemic to São Paulo (Petrongari & Simão-Bianchini 2016; Simão-Bianchini et al., 2020).

The overarching goal of this study is to test the impact of climate change in the distribution range of herbaceous and potentially invasive species of Convolvulaceae, in a region which has been disrupted by human activity and is expected to be even more so in the near future. Nine species are selected to conduct a pilot study which specifically aims to test if a duplication of CO^2^ in the atmosphere in 30-40 years time (2050-2070), will result in a decline of climatic suitability for these species.

## 2. MATERIAL AND METHODS

The state of São Paulo is located in the Southeastern Region of Brazil, extending between 19º47’ - 25º19’S latitude, and 53º06’ - 44º10’W longitude, with a surface of 248,219.627 km^2^ (IBGE, 2010). It is in a transition zone between tropical and subtropical floristic regions. According to Köppen’s climate classification (1918), São Paulo presents a total of six different climate categories: A-tropical climates, which are subdivided into 1) Af - humid tropical without a dry season; and 2) Aw - humid tropical with dry winter; and C-wet temperate climates, which are subdivided into 1) Cwa - warm with dry winter; 2) Cwb - temperate with dry winter; 3) Cfa - hot without dry season; and 4) Cfb - temperate without dry season (Köppen, 1918; Martinelli, 2010).

Of the six phytogeographic domains in Brazil (Amazon, Caatinga, Cerrado, Mata Atlântica, Pantanal, Pampas, and), only Mata Atlântica and Cerrado are represented in the State of São Paulo. However, they are very contrasting in ecological conditions, and within each of these, a range of phytophysiognomies is recognized. Cerrado is the second largest phytogeographic domain in Brazil, surpassed only by the Amazon Forest. The main phytophysiognomies in Cerrado are forest formations (*cerradão* and dry forest), savannas (*cerrado s*.*s*. and *cerrado ralo*) and *campos* (*campos sujos, campos fechados* and *campos limpos*) (Ribeiro, 1998).

Mata Atlântica phytogeographic domain is recognized for its biological diversity, and high endemism rate. Given its species richness, it is one of the areas that most requires biodiversity conservation across the planet. It is a mosaic complex with different forest formations and ecosystems, such as mangroves, sandbanks and highland fields. The high heterogeneity of the Mata Atlântica can be seen from the different environments in the same phytophysiognomy, in a landscape, or among the large ecosystems and types of vegetation. Some examples of this complex of singular heterogeneity are the Dry Forests, Dense Ombrophilous Forests, Semideciduous Forests, mangroves, restingas and *campos de altitude*, and Mata de Araucária (Cunha & Guedes, 2013).

Nine herbaceous species of Convolvulaceae, with potential weedy behaviour, were selected for this study, representing three genera - *Distimake, Evolvulus* and *Ipomoea*. These three genera have mostly a tropical distribution, with a reduced number of representatives in the subtropical areas; *Distimake* and *Evolvulus* do not occur in temperate regions. Three species from each genus were selected which were: native in the study region, taxonomically well delimited, and with a sufficiently wide geographic distribution.

A database of distribution records was built primarily from specimens with a confirmed identification by taxonomic specialists in Brazilian Convolvulaceae, with additional records extracted from the Reflora database (http://reflora.jbrj.gov.br/), where a confident species identification was possible from photographic elements. In total, 195 specimens were considered for analysis, from a total of 23 herbaria: BHCB, BOTU, CPAP, ESA, FFCL, GHSP, HRCB, IAC, K, LIL, MO, NY, PMSP, R, RB, RFA, SP, SPF, SPSF, UB, UEC, UNESP, ULM (Thiers, continuously updated) (Appendix 1).

Geographic coordinates were extracted from specimen label information; when these were not available, they were obtained through georeferencing on Google Maps (https://maps.google.com.br/) and Google Earth (http://earth.google.com/), with a careful check of the possible area of occurrence, for an estimate as accurate as possible. Databases of herbarium specimens were also consulted, such as “SPplink” (http://splink.cria.org.br/geoloc?criaLANG=pt) and “Reflora” (http://reflora.jbrj.gov.br), to search for duplicate specimen data and/or more complete geographic information, or possibly check the veracity of the obtained coordinates. The data were also revised to eliminate or correct location errors, to keep only the records with viable information and without duplication.

The potential distribution areas of the studied species were modelled in RStudio 1.3.1056 (RStudio Team, 2020) with R 3.6.3 (R Core Team, 2020) using the modleR package (Sánchez-Tapia et al., 2018). The environmental layers used were the 19 bioclimatic variables with 30 seconds resolution from the MIROC-ESM (Model for Interdisciplinary Research on Climate - Earth System Model) global climate model (GCM) available in the WorldClim database version 1.4 (Hijmans et al., 2005). After a correlation analysis processed in the modleR package, four bioclimatic variables were selected to carry out the distribution modelling: Temperature Seasonality (standard deviation ×100) (BIO4), Mean Temperature of Wettest Quarter (BIO8), Precipitation of Wettest Month (BIO13) and Precipitation of Driest Quarter (BIO17).

The species distribution modelling was performed through the Maxent algorithm for the present time (1960-1990) and projected for the future (2050-2070) with the 4.5 and 8.5 RCPs (Representative Concentration Pathways) scenarios. The final distribution models were generated through the selection of primary models with a minimum level of TSS (True Skills Statistics) equal to 0.7. The percentage of environmental suitability area for each ENM map was calculated to identify possible variations in suitability between the different ENM scenarios. The calculations were performed in the RStudio platform using the raster (Hijmans, 2020) and rgdal (Bivand et al., 2021) packages with a safe threshold of 0.5 (Franklin J, 2010; Liu, C et al.,2005). The plugin TomBiotools available for Quantum GIS 3.16.1 (QGIS Development Team, 2020) software was used to calculate the collection density and species richness through the georeferenced records dataset of the studied species, which allowed to analyse the regions where possibly there is a greater collection effort and if exists any congruence with the areas with the highest concentration of species. The record percentages of each species studied in Protected Areas (PAs) were calculated to identify the degree of species vulnerability. The procedure was carried out by crossing the collection records and the maps of the PAs present in the state of São Paulo. For this, the Point Sampling Tool plugin was used in Quantum GIS 3.16.1 to identify the georeferenced collection data belonging to the PAs and thereby calculate the percentage of these records referring to the total collections of each species.

## 3. RESULTS

The modeling of the present showed similarities in environmental suitability between *D. macrocalyx, E. glomeratus*, and *I. alba* in the northeast and central-north areas, *D. aegyptius* and *E. pterocaulon* with wide distribution and higher levels of environmental suitability in areas in the Planaltos da Bacia do Rio Paraná, *E. pusillus* and *I. carnea* subsp. *fistulosa* presenting a range of elevated areas between the Planalto Residual de Botucatu and Serra do Mar and *D. dissectus* and *I. bonariensis* with suitability areas highlighted to the east and south. The eastern region was very present in the modelling, with emphasis on Serra da Mantiqueira and Serra do Mar, including areas such as Depressão do Rio Paraíba do Sul, Planalto Residual de Botucatu, Planaltos da Bacia do Rio Paraná e the Serrania e Colinas do Ribeira.

In general, the modelling of the future showed similarities between the two levels of RCP and with the areas of suitability highlighted for the present, with specific changes for each species. Among the models of the present and future, *D. aegyptius, E. pterocaulon* and *I*.

*bonariensis* present greater stability between the suitability areas of the models, with maintenance of most of the environmental suitability between the different periods of time analysed. *D. dissectus, E. pusillus, I. alba* and *I. bonariensis* tended to concentrate their distributions in areas of suitability to the southeast in the future. The modelling of *D. macrocalyx, E. pusillus, I. alba* and *I. carnea* showed that the suitability area will tend to be lost in areas further to the north and concentrate in areas to the south, highlighting new areas such as the Planalto do Rio Paranapanema and the Planalto de Paranapiacaba *D. aegyptius, D. dissectus* and *E. pterocaulon* showed a percentage increase in the projected suitability area for the future, with emphasis on *E. pterocaulon* in the future with RCP 4.5 showing an increase of 26.73%. *D. macrocalyx, E. pusillus* and *I. alba* showed results with gain in the suitability area in the future with RCP 4.5; however, in a future scenario with RCP of 8.5, they presented a loss of environmental suitability area. *E. glomeratus, I. bonariensis* and *I. carnea* subsp. *fistulosa* were those that showed loss of area of environmental suitability in all future scenarios. Among the species that presented a decrease in areas where there is a possibility of establishment, *E. glomeratus* obtained the higher value with loss projected for the future with RCP 8.5 reaching 22.18%.

All species had a low percentage of records in PAs and, proportionally, *D. aegyptius* was the species with the most worrying results since it has the lowest number of records. *D. aegyptius* and *I. bonariensis* were the species that had the lowest and highest index of records in PAs, 6.67% and 30% respectively, with an average of 16.28% among all species. Although *D. macrocalyx* has the highest number of collection records compared to the other species, only 24.06% is present in PAs. *D. aegyptius, I. carnea, E. pusillus* and *E. glomeratus* have less than 10% of their records present in PAs.

The nine analysed species presented great variation of climatic suitability areas. *D. aegyptius* was the most widespread (77.6% of the total area), followed by *E. pterocaulon* (44.68%), *D. macrocalyx* (35.15%) and *E. glomeratus* (30.63%). The species with the most reduced modelled distribution were *I. bonariensis* (13.94%), *E. pusillus* (10.19%) and *D. dissectus* (9.03%). However, this distribution does not seem to be limited by climatic suitability, as future projections indicate that the latter are not the species experiencing the greatest decline in suitability area, but, instead, *E. glomeratus* (−22.18%), *I. carnea* subsp. *fistulosa* (−21.13%) and *D. macrocalyx* (−12.94%). Two species were hardly affected by differences in the two future scenarios (RCP 4.5 and RCP 8.5), *D. aegyptius* and *D. dissectus*,which are curiously the most widely and the least widely spread in the current modelled distributions, respectively. The remaining species generally showed a more favourable response under RCP 4.5 than RCP 8.5, with some species projected to increase in suitability area under RCP 4.5 but decline under RCP 8.5 (*D. macrocalyx, I. pusillus*, and *I. alba*); a decline in suitability area in both scenarios, more pronounced under RCP 8.5 (*E. glomeratus, I. bonariensis*, and *I. carnea*); or an increase in suitability area in both scenarios, more significant under RCP 4.5 (*E. pterocaulon*).

## 4. Discussion

Using ecological niche modelling, our study could compensate for collection biases, and infer, based on bioclimatic variables, the potential distribution of the species. For some of the species, we have discussed new predicted distribution areas for which there are no records, and which could indicate a collection bias for which some of these regions have been overlooked (e.g., areas of difficult access). Such an example is *D. aegyptius*, in which the modelled distribution suggests a strong presence in the regions of Franca and Barretos, although in our database there are no records of the species in these municipalities. The same happens with *E. pterocaulon*, which is predicted to be present in the regions of Ribeirão Preto and São José do Rio Preto; *I. alba*, in the regions adjacent to Sorocaba; *I. bonariensis* in vicinities of Bauru, and in localities close to Campinas, São Paulo and Registro; and *I. carnea* in the central regions of the state, vicinities of Bauru, and surrounding areas of Vale do Paraíba, Litoral Norte and São José dos Campos. Further fieldwork in these regions would be desirable, to confirm the presence of the species. While the state of São Paulo is widely documented and collected, species of “weeds” tend to be under collected, as seen less important or a priority for conservation, and it could still be possible that, in some regions, these species have not been collected or reported due to biases.

Of all the species included in this study, only *Ipomoea alba* L. is listed in IUCN Red List, with the category of Least Concern (Canteiro, 2021). The remaining species have a broad worldwide distribution and, although not yet assessed, are likely not under threat of extinction. The only exception is *Evolvulus pusillus*, which is endemic to Brazil and has a relatively restricted distribution in the country, in Southeast and South regions of the country.

*D. aegyptius* (a climbing herb, with great dispersion in human disturbed areas, *campos rupestres* and *cerrado*) and *E. pterocaulon* (erect herb to subshrub, predominantly found in *campos de altitude, campos limpos, campos rupestres*) have demonstrated a positive response to climate change, even in the most extreme scenarios, which is an indication that they may require weed management in the future, as they will progressively occupy new environments.

*D. dissectus* (a climber, occupying a range of vegetation types, predominantly environments with high level of humidity) presents a slightly projected increase in climatic suitability range, and it may not represent a significant threat to ecosystem management in the future, but considering its positive response to climate change, it should still be tracked for any possible expansion of its distribution range with undesired consequences in ecosystem balance.

*E. glomeratus* (erect herb to subshrub), *I. bonariensis* (climber) and *I. carnea* subsp. *fistulosa* (climber) are the species that will be most negatively impacted by climatic change, with their climatically suitable range suffering significant declines, especially in the most extreme climate change scenario (RCP 8.5). Considering that they are widespread species across tropical regions, this may not represent a threat to the conservation of the species, but it could have a significant impact in the floristic diversity of the state of São Paulo, with consequences also for the local fauna (e.g. pollinators).

The average among all species presented only 16.28%, which may indicate the poor positioning of the reserves in relation to the species. With a 30% index of records in PAs, *I. bonariensis* as well as *E. pterocau*lon does not present, among its main phytophysiognomies, the Anthropic Area, being present in areas that possibly have PAs. Second, with 29.63% is *D. dissectus*, which covers a larger area in relation to phytophysiognomies where it occurs. The number of records in PAs is still low, ecological and systematic surveys are suggested to expand understanding about the distribution of species and provide a rigorous assessment of the conservation status of each species, regardless of the location of registration sites in relation to PAs. Global climate change and invasive species can also affect species within reserves and the potential impacts of both events need to be better understood (Lemckert, F. et al., 2009).

Understanding how weeds will be affected by climate change is important for ecosystem management, as much as studying species that are at greater risk of extinction. In the absence of key climatic modelling studies for Convolvulaceae, this study hopes to be a useful tool and reference for understanding climatic suitability in the present as well as future climatic conditions, in this important but ecologically overlooked taxonomic group for the Brazilian flora, and which contributes significantly to the floristic composition of the Cerrado and Mata Atlântica phytogeographic domains. It is increasingly important to conduct such modelling studies in very urbanized areas such as the state of São Paulo, where there are several habitat threats and natural ecosystems that need to be carefully managed.

Hence, conservation action in PAs and the areas here identified as climatic refuges, will be key to avoid worsening the environmental threats in the state, and mitigate potential loss in floristic diversity. Herbaceous species, such as most Convolvulaceae, are often neglected in general collections. The results of this study demonstrate that the potential area of distribution of a herbaceous species can be much wider than its current distribution, based on observations and collections, suggest. This further highlights the importance of using modelling tools to fill in collection gaps.

## 4. ACKNOWLEDGEMENTS

The first author thanks CNPq for the grant offered to develop this project (Bolsa PIBIC 137929/2017-0), and all the researchers, curators and students at Instituto de Pesquisas Ambientais, São Paulo (Herbarium SP) for all the support and infrastructure.

In particular, the authors would like to thank Marcela Inácio da Silva and Sónia Aragaki (Instituto de Pesquisas Ambientais), for useful comments on the manuscript.

Ana Rita Simões would like to thank CAPES (Brazil) for funding her research (2015-2018), Grant BJT 88881.067993/2014-01.

For the photographs, we thank Mayara Pastore, Diego Santos, Henrique Moreira, Fernanda S. Petrongari, Ulisses Fernandes.

